# Antagonism among DUX family members evolved from an ancestral toxic single homeodomain protein

**DOI:** 10.1101/2023.01.21.524976

**Authors:** Darko Bosnakovski, Erik A. Toso, Elizabeth T. Ener, Micah D. Gearhart, Lulu Yin, Felipe F. Lüttmann, Alessandro Magli, Ke Shi, Johnny Kim, Hideki Aihara, Michael Kyba

## Abstract

Double homeobox (DUX) genes are unique to eutherian mammals and normally expressed transiently during zygotic genome activation. The canonical member, DUX4, is involved in facioscapulohumeral muscular dystrophy (FSHD) and cancer, when misexpressed in other contexts. We evaluate the 3 human DUX genes and the ancestral single homeobox gene sDUX from the non-eutherian mammal, platypus, and find that DUX4 activities are not shared with DUXA or DUXB, which lack transcriptional activation potential, but surprisingly are shared with platypus sDUX. In human myoblasts, platypus sDUX drives cytotoxicity, inhibits myogenesis, and induces DUX4 target genes, particularly those associated with zygotic genome activation (ZGA), by binding DNA as a homodimer in a way that overlaps the DUX4 homeodomain crystal structure. DUXA lacks transcriptional activity but has DNA-binding and chromatin accessibility overlap with DUX4 and sDUX, including on ZGA genes and LTR elements, and can actually be converted into a DUX4-like cytotoxic factor by fusion to a synthetic transactivation domain. DUXA competition antagonizes the activity of DUX4 on its target genes, including in FSHD patient cells. Since DUXA is an early DUX4 target gene, this activity potentiates feedback inhibition, constraining the window of DUX4 activity. The DUX gene family therefore comprises cross-regulating members of opposing function, with implications for their roles in ZGA, FSHD, and cancer.

**HIGHLIGHTS:**

- Platypus sDUX is toxic and inhibits myogenic differentiation.
- DUXA targets overlap substantially with those of DUX4.
- DUXA fused to a synthetic transactivation domain acquires DUX4-like toxicity.
- DUXA behaves as a competitive inhibitor of DUX4.

## INTRODUCTION

The double homeobox gene (DUX) family is unique to eutherian mammals ^1^. Although some homeodomain-containing transcription factors have a second DNA-binding domain of a different type, the presence of two homeodomains in tandem does not occur elsewhere in the tree of life. The DUX gene family is derived from an ancestral single homeobox gene, named *sDUX*, present in fish, lizards, birds and non-eutherian mammals, via tandem duplication of the *sDUX* single homeobox ^2^. This new double homeobox gene then expanded, diverged within the ancestral placental mammal genome into three clades: DUXA, DUXB and DUXC, with DUXC genes uniquely expanding massively into long tandem arrays, while DUXA and DUXB remained present at low or single copy ^2^ (Fig. 1a). There has been significant sequence diversion within each homeodomain (HD), demonstrated by the fact that sequence dendrograms do not cluster HD1 sequences from DUXA, B, and C together, nor those of HD2, but rather, each HD of each DUX clade is rooted deeply, each practically as distant from the sDUX HD as the others (Fig. 1b). In humans, the DUXC representative is named *DUX4* ^3^, and its tandem arrays are subject to repeat-induced silencing ^4^. If the array contracts to low copy numbers, the silencing becomes inefficient, leading, by way of leaky expression of *DUX4* ^5^, to the disease, facioscapulohumeral muscular dystrophy (FSHD) ^6^. DUX4 is toxic to myoblasts and other cell types ^7,8^, and also impairs myoblast differentiation when expressed at low non-toxic levels ^8,9^. Besides the homeodomains, DUXC family members share a conserved domain at the C-terminus, which is not found in DUXA and DUXB family members ^1,2^. The DUX4 C-terminal domain is an activation domain necessary for target gene expression ^10,11^, is essential for cytotoxicity ^12^, and confers transcriptional activation and pioneer activity to DUX4 by interacting with histone acetyltransferases p300 and CBP to mediate histone H3 acetylation, chromatin opening and transcription of target genes ^13^. This C-terminal conservation is not found in various sDUX representatives, thus it is unclear whether sDUX is a transcriptional activator.

**Figure 1.**
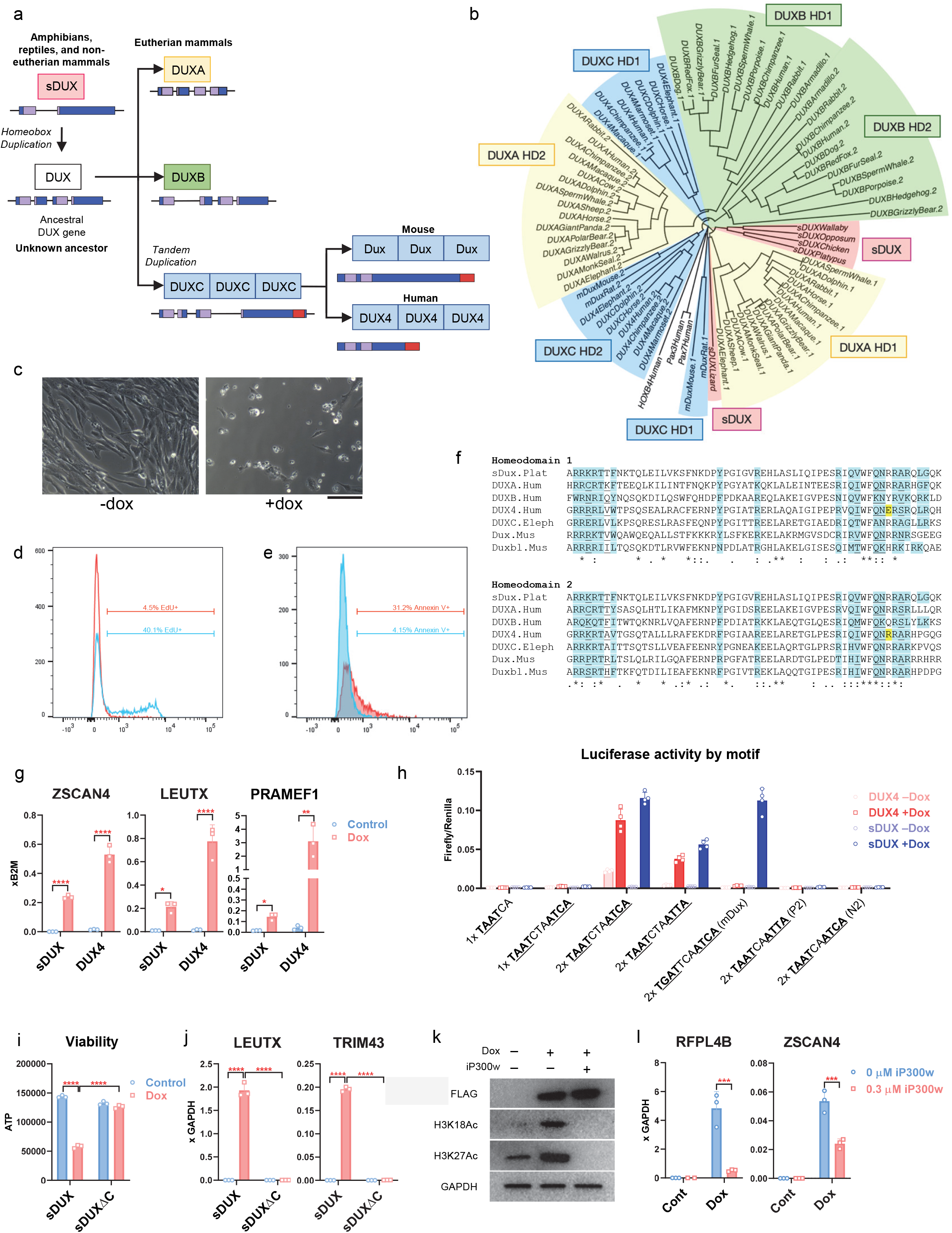
Platypus sDUX displays functional overlap with DUX4. a. Evolution of the DUX gene family from an ancestral single homeobox gene in the common ancestor to eutherian mammals. b. Sequence dendrogram of DUX homeodomains. DUX family members are shaded yellow (DUXA), green (DUXB), and blue (DUXC); and sDUX is shaded red. c. Morphology of LHCN-M2isDUX human myoblasts in the absence (left) or presence of 100 ng/mL dox for 4 days to induce sDUX (right). Scale bar, 100 um. d. EdU incorporation in LHCN-M2isDUX human myoblasts after 6 hrs in the absence (blue) or presence (red) of 500 ng/mL dox to induce sDUX. e. Annexin V incorporation in LHCN-M2isDUX human myoblasts after 24 hrs in the absence (blue) or presence (red) of 500 ng/mL dox to induce sDUX. f. Sequence comparison of the sDUX homeodomain with HD1 and HD2 of the 3 human DUX family members, DUXC from elephant and mouse, and mouse DUXBL. Known or predicted DNA binding regions are highlighted in blue, and residues known or predicted to have specific nucleotide base contacts are underlined. Yellow shading indicates the E and R residues responsible for TAAT or TGAT specificities of DUX4 HD1 and HD2, resp. Asterisks indicate positions with a conserved residue, colons indicate conservation of strongly similar biochemical properties, periods indicate conservation between amino acids of weakly similar properties. g. RTqPCR for *ZSCAN4, LEUTX* and *PRAMEF1* in LHCN-M2iDUX4 parent and LHCN-M2-isDUX myoblasts induced with 200 ng/mL doxycycline for 14 hours. Data are presented as mean ± SEM; *p<0.05, ***p<0.01**, ****p<0.0001 by one-way ANOVA, n=3. h. Luciferase assays in comparing activity of DUX4 and sDUX on different versions of the double homeodomain motif in 293T cells exposed to 500 ng/mL dox for 24 hrs. The core HD binding sites are underlined. DUX4 and DUX cores are separated by 3 nt while the PAX7 recognition motif cores are separated by 2 nt. Copy number of motifs indicated by 1x or 2x. i. Viability assay for 293T cells after 48 hrs of expression (100 ng/mL dox) of full length sDUX *vs*. sDUXΔC, lacking the C-terminal 74 amino acids. j. RTqPCR for DUX4 target genes *LEUTX* and *TRIM43* after expression of sDUX or sDUXΔC in the same cells in panel I (500 ng/mL, 24 hrs). k. Representative western blots with acetylation specific-antibodies in human myoblasts expressing sDUX and treated with the p300/CBP-specific HAT inhibitor, iP300w. LHCN-isDUX cells were induced with 200 ng/ml doxycycline and concurrently treated with 0.3 μM iP300w for 12 hours. l. RTqPCR for sDUX target genes *RFPL4B* and *ZSCAN4* in the presence or absence of iP300w on the cell presented in panel k.

Both the mouse and human DUXC genes (*Dux* and *DUX4*) are briefly expressed in the cleavage stage embryo, at the 2- and 4-cell stage, respectively, where they activate a subset of genes expressed during zygotic genome activation (ZGA) ^14,15^, and so were proposed to be key inducers of ZGA ^16^. However, because most species lack DUXC genes, because mouse *Dux* array knockouts are viable ^17-19^, because the majority of ZGA-associated genes are still expressed in the absence of *Dux*, because the rate of failure of KO embryos to develop to blastocyst *in vitro* is much higher than their rate of failure *in vivo* ^18^, and because growth defects characterize a subset of *Dux* KO embryos, it has been proposed that this transient expression and downstream 4/8 cell target gene expression primes the embryo for later efficacious placentation rather than initiating ZGA *per se* ^18^. In any case, the expression of DUXC family members is tightly restricted to early blastomere stages of embryogenesis, with a strong need for silencing in later somatic cells.

DUX genes have also been implicated in cancer, with one study suggesting that DUX4 expression leads to downregulation of MHC proteins and thereby poor outcomes ^20^, and another showing that amplification of murine *Duxbl* cooperates with p53 inactivation to drive rhabdomyosarcoma ^21^. Interestingly, the latter study found that in human cells, *DUXA* was a strongly induced downstream target of DUX4 (*DUXB* was also induced at a lower level). This suggested the possibility of feed-forward regulation within the family. Because DUX4 is virtually impossible to detect at the protein level in FSHD patient muscle biopsies, it was suggested that early transient DUX4 expression may set up a pathological program that persists long after a DUX4 burst ^22^. Recent RNA-seq data on cultured FSHD myoblasts revealed a population of *DUXA*+ cells that lacks *DUX4* expression, prompting a model in which DUXA takes over the induction of DUX4 target genes after a burst of DUX4 expression ^23^, however the transcriptional activity of DUXA is unknown. A mechanistic understanding of the role of the DUX family in development, FSHD and cancer, thus requires understanding the interplay between DUXA, B and C and how their functions evolved from and relate to that of the ancestor, sDUX.

## RESULTS

### sDUX binds DUX motifs and is toxic to human myoblasts

To frame and root the function of the DUX family, we began by investigating sDUX from a species closely related to eutherian mammals, namely the monotreme *Ornithorhynchus anatinus* (platypus). Remarkably, doxycycline (dox)-inducible expression of platypus sDUX led to cell death of human myoblasts, in a manner very similar to that seen with DUX4 (Fig. 1c). sDUX-expressing myoblasts ceased proliferating and underwent apoptosis as assessed by EdU incorporation (Fig. 1d) and annexin V staining (Fig. 1e). Structural studies on the DUX4 homeodomains ^24^ have identified a critical residue (E vs R) in helix 3 that specifies the TAAT vs. TGAT core preference (yellow residues in Fig. 1f). DUX4 HD1, which bears E at this site, prefers TAAT, and HD2, which bears R, prefers TGAT (ATCA in the motif, since the HDs bind head-to-head), while both murine DUX HD1 and HD2 bear R and thus prefer TGAT (ATCA at the 3’ end of the motif). The sequence of the sDUX homeodomain predicts a DNA-binding specificity similar to that of murine DUX. Despite this prediction, RTqPCR revealed that in human myoblasts, sDUX induced the canonical DUX4 target genes *ZSCAN4, LEUTX*, and *MBD3L2* (Fig. 1g). To probe the sequence specificity of sDUX, we tested the ability of sDUX and DUX4 to transactivate from variations of double homeodomain recognition motifs on luciferase expression plasmids bearing two copies of each motif upstream of a minimal promoter. While DUX4 clearly preferred the non-palindromic TAAT---ATCA motif (named N3 for non-palindromic with 3 nt spacing), sDUX was able to drive luciferase expression from both the DUX4 N3 motif and the murine DUX-type palindromic TGAT---ATCA P3, as well as from the palindromic TAAT---ATTA P3 motif, suggesting a broader DNA-binding spectrum for the ancestral protein compared to individual DUXC representatives from human and mouse (Fig. 1h). Half-motif constructs (TAAT or TGAT alone) showed no activity, suggesting that sDUX binds cooperatively as a dimer to motifs with two core HD recognition sites. Furthermore, transactivation was detectable only at extremely low levels when motifs were present in a single copy, indicating cooperativity of transactivation when multiple binding sites regulate a given promoter. In view of the sequence similarity between DUX and PAX HDs, and a competitive interaction that has been described between DUX4 and PAX7 ^8^, and recent observations of a PAX7 profile that has been detected in FSHD muscle biopsy RNA-seq ^25,26^, we also tested the Pax7 motif, which has TAAT cores but spaced by 2 nt, as well as variants in which one or both cores were TGAT. sDUX, like DUX4, had no activity on HD core motifs spaced by 2 nt.

As the activation domain of DUX4 is C-terminal, we tested the activity of an sDUX mutant lacking the C-terminal 74 amino acids. sDUXΔC was both non-toxic (Fig. 1i) and unable to induce target genes (Fig. 1j) indicating that its activation domain is localized within its C-terminus and is necessary for cytotoxicity, like DUX4 ^8,11^. Because the C-termini of DUX4 and sDUX do not show obvious sequence conservation, we tested their mechanistic similarity. One of the most unusual features of the DUX4 C-terminus is that its expression leads to a global increase in nuclear H3K18 and H3K27 acetylation ^13,27^. We found that cells expressing sDUX likewise displayed rapid and massive increases total H3K18Ac and H3K27Ac (Fig. 1k). The DUX4 activation domain is exquisitely dependent on the histone acetyltransferase (HAT) activity of p300 and CBP ^27,28^. We found that H3 acetylation driven by sDUX was reversed by treatment with iP300w (Fig. 1k), a specific inhibitor of the HAT activity of p300 and CBP, as was the case for DUX4 ^27^. Likewise, transcriptional activation by sDUX was also suppressed by iP300w (Fig. 1l), indicating that the activation domain of sDUX requires a p300/CBP cofactor, like that of DUX4.

### Structural basis of sDUX DNA binding

To understand how sDUX could bind DUX4 targets when it has only a single HD, while DUX4 has two HDs of different core specificity, we tested the ability of bacterially produced sDUX protein to bind to double-stranded DNA bearing predicted recognition motifs using the electrophoretic mobility shift assay (EMSA). sDUX generated a pattern of band-shifts similar to that of DUX4 on both non-palindromic (TAAT---ATCA) and palindromic (TGAT---ATCA or TAAT---ATTA) motifs (Fig. 2a). Although a weak band corresponding to sDUX monomer bound to DNA was observed at lower protein concentrations, sDUX bound to these DNA motifs primarily as a homodimer, suggesting high cooperativity. In contrast to the luciferase transcriptional assay, sDUX was also able to bind as a monomer to DNA substrates bearing either TAAT or TGAT half-motif, for which DUX4 shows no binding (Fig. 2b). Similarly, sDUX bound as a monomer to the substrate with 2 nt separating TGAT and ATCA, incorrect spacing for DUX4 (Fig. 2b,c). These results corroborate the results of the luciferase reporter assay that sDUX has a broader target DNA sequence selectivity, although they suggest that not all sites bound by sDUX in vitro (e.g., monomer sites) are transcriptionally active.

**Figure 2.**
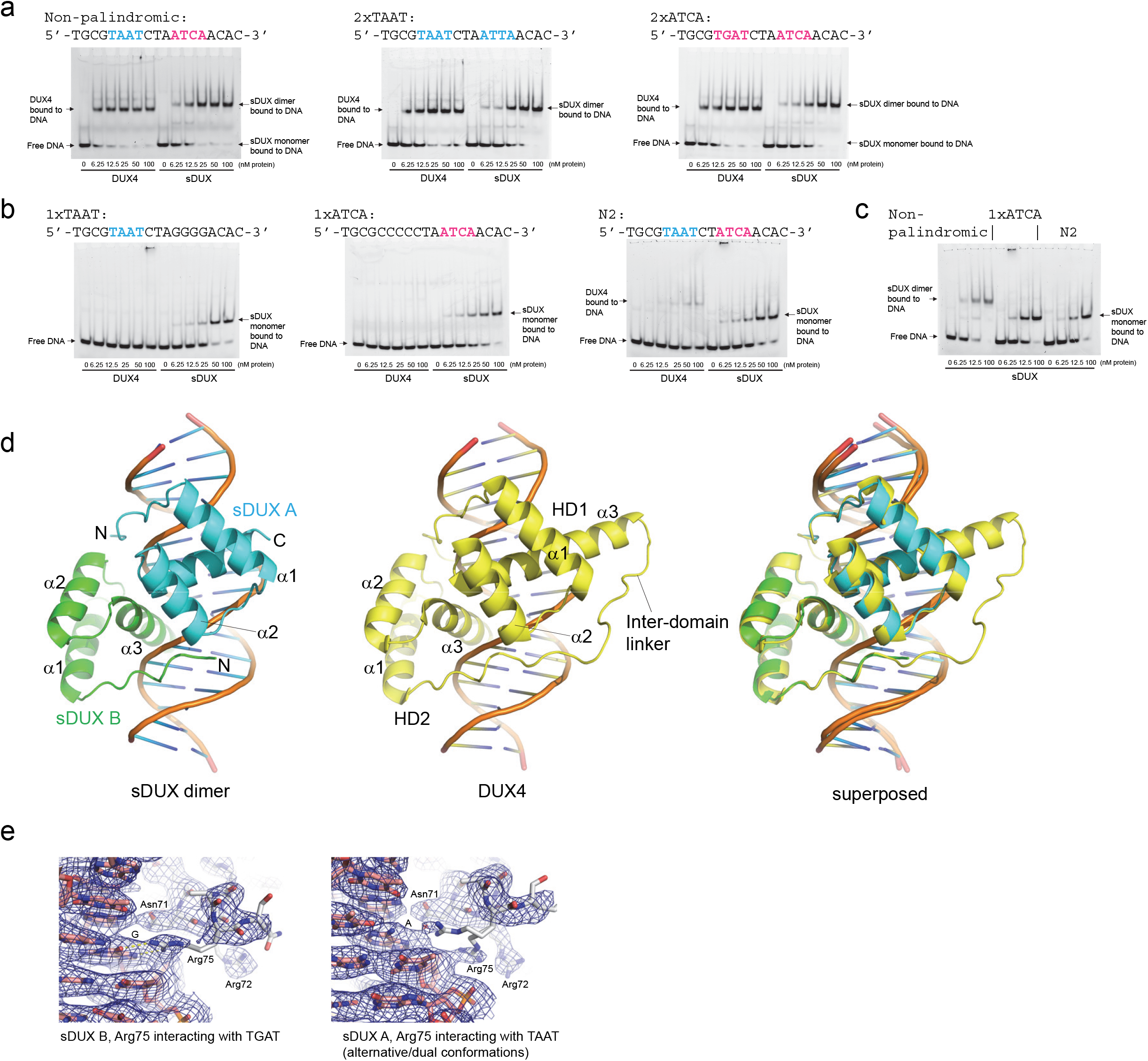
Structural basis of sDUX interaction with DNA. a. Binding of DUX4 HD1-HD2 and sDUX HD to the non-palindromic (TAAT---ATCA) and palindromic (TGAT---ATCA or TAAT---ATTA) motifs tested in EMSA. All DNA substrates were double-stranded DNA. Sequence of the fluorescently labeled strand is shown above each gel. b. Binding of DUX4 HD1-HD2 and sDUX HD to the TAAT and TGAT half-motifs or a non-palindromic motif with 2 nt spacing (N2) tested in EMSA. c. Binding of sDUX HD to the three different types of DNA substrates, highlighting distinct mobilities of dimeric vs. monomeric complex in EMSA. d. Crystal structure of sDUX homodimer bound to the non-palindromic target motif with 3 nt spacing, the structure of DUX4 double homeodomain with the same DNA substrate as reported previously ^24^, and their superposition. e. Differential interaction of sDUX Arg75 with the TAAT and TGAT motifs in the crystal structure. 2Fo-Fc electron density for the structure with Br-dU-containing DNA is shown as blue mesh, contoured at 0.9 σ.

We then determined the crystal structure of the sDUX homeodomain in complex with the non-palindromic DNA substrate that had previously been co-crystallized with the DUX4 double homeodomain (Supplementary Table 1) ^24^. As expected from the biochemical observations above, sDUX binds to this target sequence as a head-to-head homodimer (Fig. 2d). Except for the interdomain linker and the extended α-helix 3 of HD1, which are unique features for DUX4, the sDUX-DNA and DUX4-DNA complexes show overall very similar structures with a root mean square deviation (r.m.s.d.) of 0.75 Å for the protein backbone and all DNA atoms (Fig. 2d). The two sDUX homeodomains interacting with the TAAT and TGAT half-sites also show high structural similarity, although they differ in the conformation of Arg75 from a characteristic basic patch near the C-terminus of α-helix 3 (^72^RRAR^75^; Fig. 1f). The side chain of Arg75 forms bidentate hydrogen bonds with the guanine base of 5′-TGAT-3′ in the major groove, while it shows alternative positioning in interacting with the 5′-TAAT-3′ motif (Fig. 2e). This is reminiscent of the differential positioning of the corresponding Arg residues of DUX4 (Arg73 in HD1 and Arg148 in HD2) in the DUX4-DNA complex, which we previously showed is caused by a difference in key upstream residues (Glu70 in HD1 *vs*. Arg145 in HD2) ^24^.

Collectively, these data emonstrate that sDUX shows remarkable similarities to DUX4, including its primary architecture, mode of DNA-binding, toxicity, DUX4 target gene expression, and p300-dependence, suggesting that the DUXC clade of the DUX family has retained the primary molecular properties of the ancestral gene at the root of the family.

### DUXA and DUXB lack significant transcriptional activation potential

To understand the relationship of the A and B branches vis-à-vis C of the DUX family to sDUX, we next expressed the human DUXA and B proteins in human myoblasts, in their native form; and because their transactivation potential was uncertain due to the lack C-terminal sequence conservation with the DUX4 transcriptional activation domain we also fused them to a C-terminal VP64 activation domain ^29^; and we compared them to human DUXC (i.e., DUX4), and to sDUX. Neither DUXA nor DUXB displayed any toxicity. Interestingly however, the DUXA-VP64 fusion was toxic to human myoblasts, while DUXB-VP64 was not (Fig. 3a). We then evaluated the ability of these constructs to activate expression using luciferase reporters bearing two copies of the 3 flavors of DUX motif, as well as the Pax7 (P2 bearing 2 nt spacing) motif (Fig. 3b, Supplementary Fig. 1). DUXA and DUXB were both transcriptionally inactive in this assay, although the fusion to VP64 converted DUXA into an activator, and revealed that DUXA recognizes both the P3 TGAT-- -ATCA murine DUX motif and the N3 TAAT---ATCA DUX4 motif equally well, and also, but more weakly, the P3 ATTA---ATTA motif, behaving very like sDUX in this assay. To evaluate the ability to induce DUX4 target genes, we performed RNA-seq on DUXA, B, VP64 derivatives and sDUX cells exposed to dox for 6 hours, and unmodified cells and compared to previously published data for DUX4 in the same system ^13^. DUXA, DUXB, and DUXB-VP64 showed no induction of DUX4 target genes, whereas DUXA-VP64 and sDUX both showed a pattern of upregulation of these genes (Fig. 3c). These data strongly suggested that sDUX and DUXA may bind to many DUX4 target sites. DUXA apparently lacks a transcriptional activation domain of its own but acquires transactivation potential if provided with a heterologous activation domain at its C-terminus.

**Figure 3.**
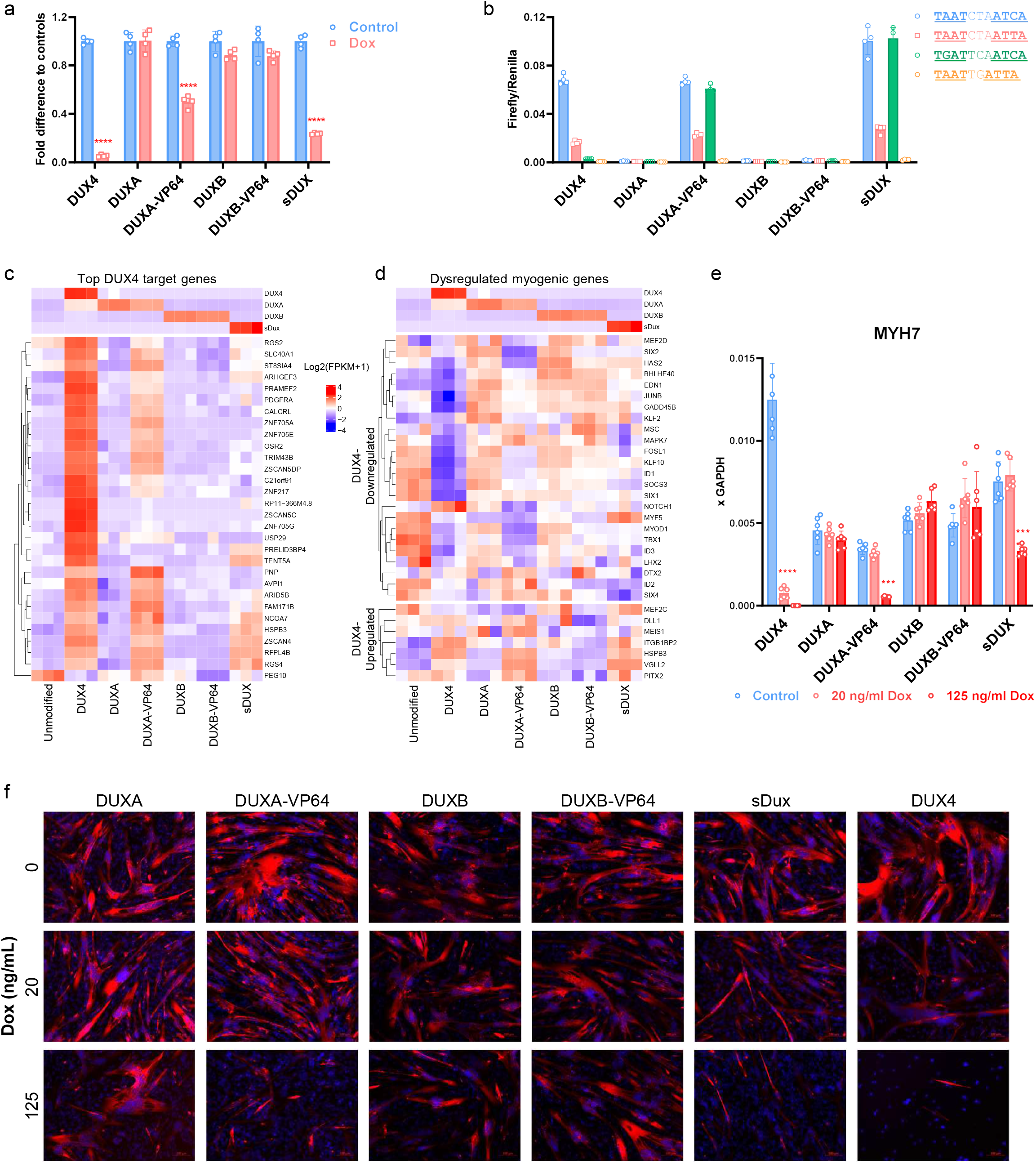
Toxicity and myogenic differentiation effects of DUX family members and VP64-fusions. a. Viability assay in LHCN-M2 human myoblasts after 96 hrs of expression of different DUX family proteins induced with 200 ng/mL doxycycline. b. Luciferase assays in 293T cells induced with 1 μg/mL dox for 24 hrs to express various DUX family proteins and VP64 derivatives on the 3 versions of DUX motif (N3 and P3) as well as a motif with spacing of 2 nt (P2-type). c. Heat map of the top DUX4 target genes showing expression in cells expressing various DUX family members and VP64 derivatives after 6 hours of induction (200 ng/mL dox). d. Heat map of the top myogenic genes affected by DUX4, and the effect of other DUX family members, and DUXA- and DUXB-VP64 activation domain fusions on this set. Genes downregulated by DUX4 shown above; genes upregulated by DUX4 shown below. Cells were induced by 200 ng/mL dox for 12 hours. e. RTqPCR for MYH7, a marker of differentiation, after differentiation with 72 hrs of expression of different DUX family proteins at 20 or 125 ng/mL dox. f. Representative immunostaining for sarcomeric myosin heavy chain on the same cells shown in panel e.

### DUXA-VP64 interferes with myogenesis

One of the hallmark activities of DUX4 is inhibition of myogenesis ^8^, and this is associated with its ability to perturb (primarily by downregulation) myogenic gene expression ^9^. We therefore evaluated the set of myogenic genes perturbed by DUX4 ^9^ within the profiles of each DUX family member (Fig. 3d). While native DUXA and DUXB had virtually no effect on these genes, sDUX and DUXA-VP64 showed DUX4-like perturbations of many of these genes, including the key master regulator, *MYOD1*, which was strongly downregulated by both factors. This again indicates the similarity of sDUX to DUX4 and further suggests that DUXA may have a biologically meaningful degree of overlap in DNA-binding with DUX4, revealed here by converting it into an activator by fusion to VP64, whereas DUXB does not. To test the phenotypic effect of these myogenic changes, we subjected the cell lines to differentiation in the presence of doxycycline. DUXA-VP64 and sDUX both showed suppression of *MYH7*, a differentiation marker, as well as of morphological differentiation to multinucleated pan-myosin heavy chain+ myotubes, similar to DUX4 (Fig. 3e,f).

### Transcription and chromatin profiling reveals relationships among DUX family proteins

In order to definitively evaluate the regulatory potential of DUX family members, we performed RNA-seq in both the presence and absence of a 12-hr pulse of dox on DUX4, DUXA, DUXB, DUXA-VP64, and sDUX. Principal component analysis showed transcriptional profiles of all cell lines in the absence of dox to be clustered, with the exception of DUX4, which showed a leftward shift on PC1, the axis that contains 62% of variance amongst all conditions (Fig. 4a). Dox induction led to a much greater left shift for DUX4, indicating that the leftward shift of the control reflects very low background expression of DUX4 targets in the absence of dox. In the presence of dox, both DUXA and DUXB remained close to the cluster containing the negative controls, supporting the notion that they are transcriptionally inactive. Both sDUX and DUXA-VP64 showed a degree of similarity to DUX4 in the form of a slight leftward shift on PC1 (where most variance is contained), but also an upward shift on PC2 (which contains only 8% of the variance). This indicates that there are unique aspects of the sDUX profile that DUXA-VP64 participates in but DUX4 does not. The relationships between the expression profiles of sDUX, DUX4, and DUXA-VP64 are striking (Fig. 4b) with significant overlapping gene sets (Fig. 4c). A large fraction of the genes induced by each factor in human myoblasts comprises genes expressed up to and during the human 8 cell stage ^30^, here referred to as ZGA genes (sDUX, 119 of 671, 18%; DUXA-VP64, 101 of 629, 16%; DUX4 436 of 2523, 17%). Within the ZGA gene set, there is slightly larger overlap, for example 74% of the sDUX ZGA targets are co-regulated by DUX4 (*vs*. 64% of all targets) as are 70% of the DUXA-VP64 ZGA targets (*vs*. 45% of all targets). Repetitive elements, especially LTR-like repeats, are prominently and specifically expressed in early cleavage stages ^31^, and represent a significant set of ZGA-related targets of DUX4 and mouse DUX. We found that both sDUX and DUXA-VP64 induced transcription of repetitive elements, with LTR-type elements being most abundant among these, and sDUX sharing with DUX4 the ability to induce LINE elements (Fig. 4d). Strong correlations were seen among the transcriptional profiles of sDUX, DUX4, and DUXA-VP64 in both conventional genes (Fig. 4e, upper left) and repetitive elements (Fig. 4e, lower right); with the greatest correlations being between sDUX and DUX4 (Fig. 4f,g).

**Figure 4.**
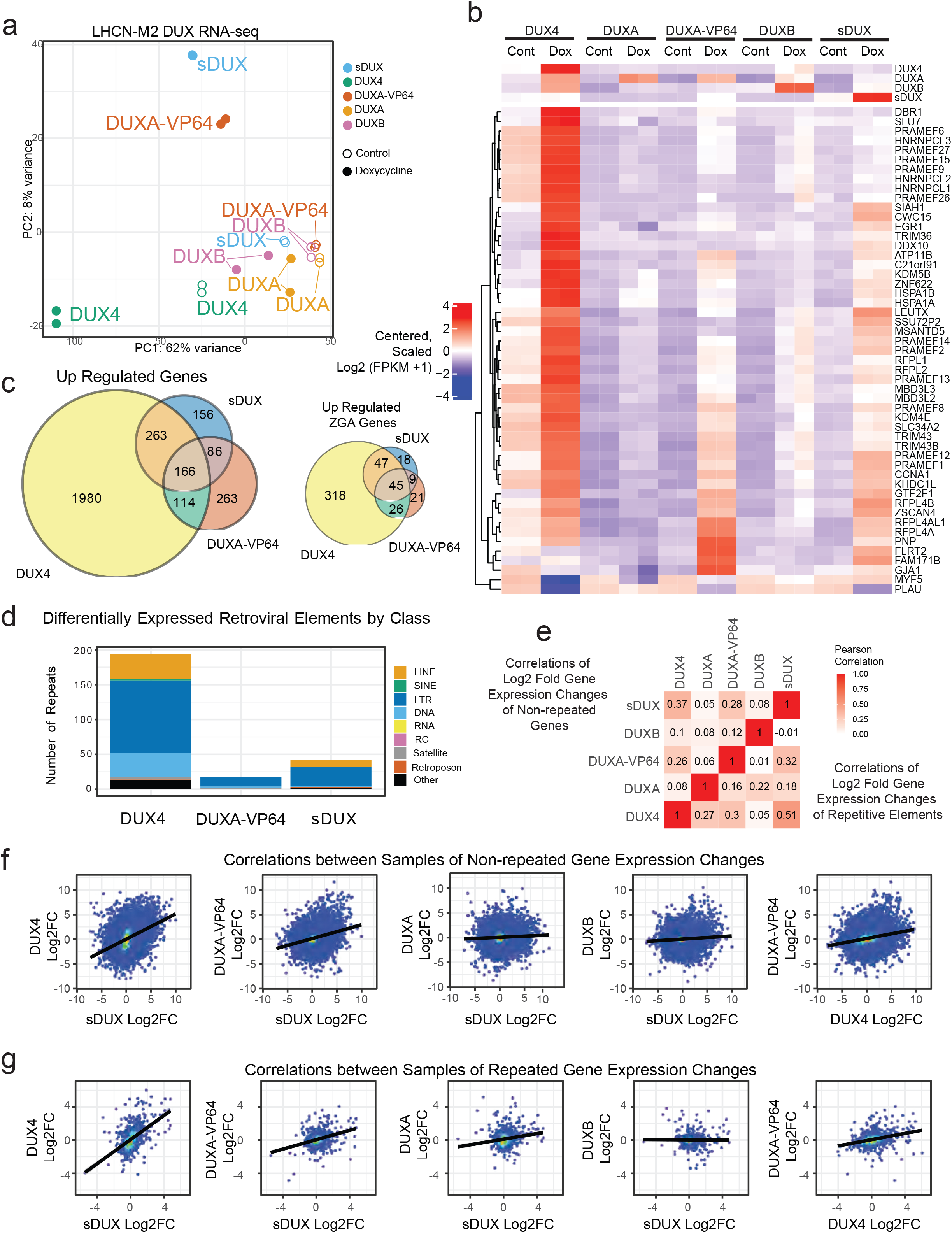
Transcriptional profiles of DUX family members. a. Principal component analysis of gene expression profiles in the absence (open circles) and presence (closed circles) of 250 uM dox for 12 hrs. DUX4 expressing cells are strongly shifted along PC1 which accounts for 62% of the variance while DUXA-VP64 and sDUX expressing cells are more prominently shifted along PC2 which contains 8% of the variance. b. Heatmap representation of log2 transformed FPKM values for the top 50 differentially expressed genes among all factors. Expression of *DUX4, DUXA, DUXB* and *sDUX* is shown in the upper panel. c. Venn diagrams showing overlap in upregulated gene sets among total genes (left) and ZGA genes (right). Upregulated genes for each set were defined as having a greater than 2-fold change, an Benjamini-Hochberg adjusted p-value less than 0.05 and a mean FPKM value > 2.5 in the dox induced sample. d. Distribution of repeat classification among the differentially expressed retroviral elements for DUX4, DUXA-VP64 and sDUX. Differentially expressed elements were defined as being upregulated or downregulated greater than 2-fold and having a Benjamini-Hochberg adjusted p-value less the 0.05. e. Pearson correlations of log2 fold changes in expression upon induction with dox between each of the DUX factor. The upper left half of the matrix corresponds to conventional genes whereas the lower right half of the matrix corresponds to log2 fold changes for repetitive elements. f. Scatter plots of the log2 fold changes in conventional expression between DUX factors. Points representing individual genes are colored according to density to show the higher density of points near the center. A linear fit of each correlation is shown as a black line. g. Scatter plots similar to panel f for repetitive elements.

We next investigated global chromatin changes with ATAC-seq on the same set of cells after 12 hours of dox exposure to induce expression of DUX proteins. ATAC-seq peak profiles for the control untreated cells clustered together, and again, major differences after dox induction were observed only with DUX4, sDUX, and DUXA-VP64, while DUXA and DUXB continued to cluster with the controls, although DUXA showed a greater change in response to dox than DUXB (Fig. 5a). Numerous co-localized dox-induced peaks near ZGA genes were found, for example *ZSCAN4* and *PRAMEF12*, in which DUX4, sDUX, DUXA-VP64, as well as DUXA peaks overlapped (Fig. 5b,c).

**Figure 5.**
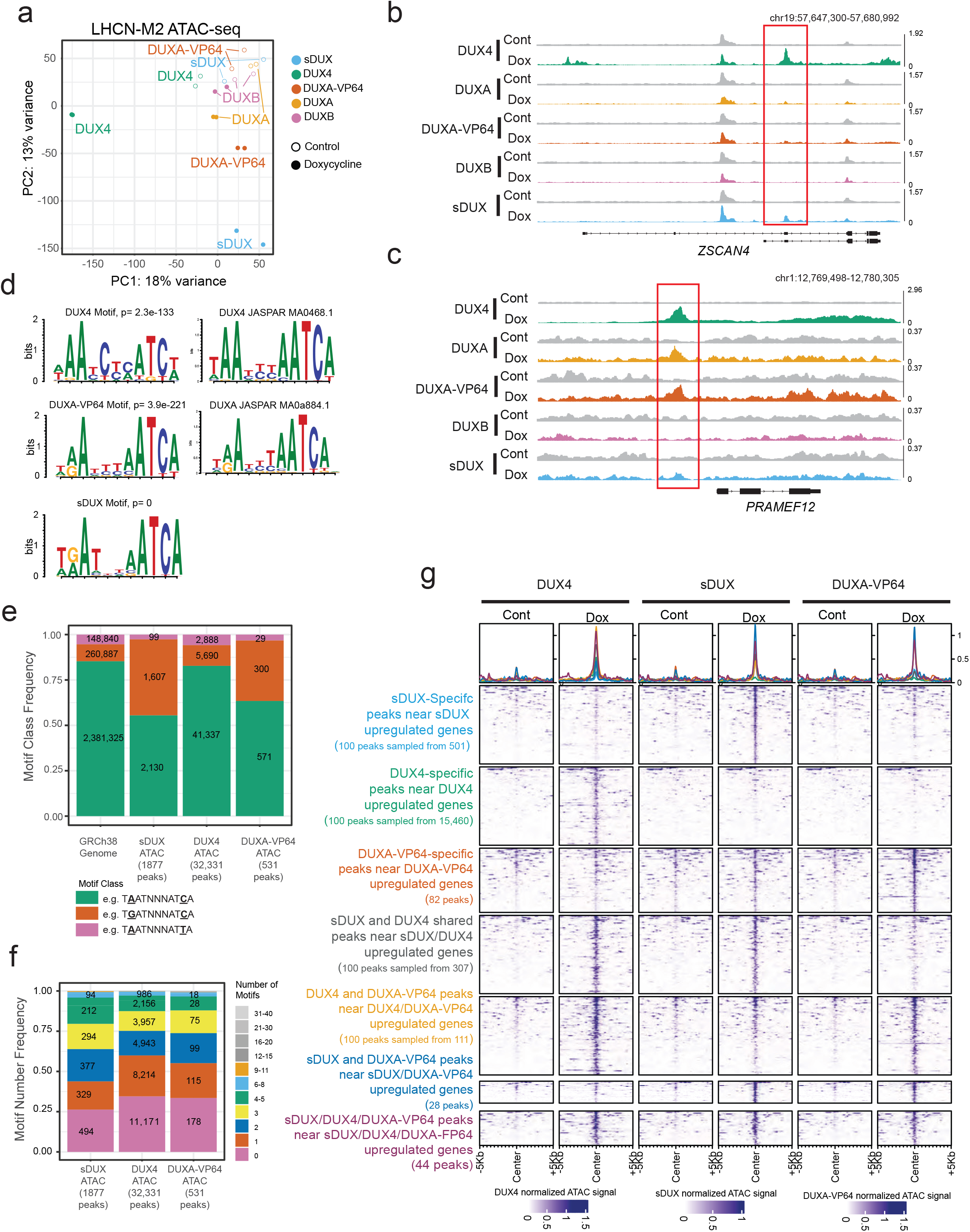
Chromatin changes induced by DUX family members. a. Principal component analysis of ATAC-seq data in the absence (open circles) and presence (closed circles) of 250 uM dox for 12 hrs. Called peaks from each factor were combined into a composite set of genomic locations that had changes in DNA accessibility in one or more samples. A matrix of the number of reads within each region across all the samples was used to calculate the principal components. b. ATAC-seq coverage tracks for each DUX factor at the *ZSCAN4* locus in the absence (gray) or presence (colored according to panel a) of dox. Each track shown is a combination of two biological replicates normalized to the total number of reads per million. The scale for each pair of tracks is shown at the right and the gene structures is shown below. The red box corresponds to peaks that appear for some DUX factors upon the addition of dox. c. ATAC-seq coverage similar to panel B for the *PRAMEF12* locus. d. The DNA sequences from the top 500 peaks were used to define de novo motifs for DUX4, DUXA-VP64 and sDUX. The top motif and its p-value are shown. The DUX4 and DUXA-VP64 motifs are similar to the previously defined motifs for DUX4 and DUXA repectively (https://jaspar.genereg.net/). e. The de novo motifs identified from the top 500 DUX4, DUXA-VP64 and sDUX peaks were identified across each dataset using a 95% scoring threshold. Classification of each sequence is based on the nucleotide at the 2nd and 10th position. f. The number of sequence motifs identified in panel e per ATAC-seq peak were counted to show that peaks frequently contain multiple copies of the DUX motif in all three datasets. g. Heatmaps of ATAC-seq coverage for peaks within 10 kb of an upregulated gene. Peaks were classified into one of seven groups depending on whether they were unique to a particular dataset or shared between two or more datasets. Groups that contained more than 100 peaks were randomly subsampled to 100 peaks. Coverage was normalized to total counts per million reads of each dataset and then plotted on the scale defined by the dox induced dataset shown below the heatmaps.

Interrogating peaks for the presence of overrepresented sequence motifs revealed variants of the 11 bp motifs previously associated with DUX4 ^10^ and mouse DUX ^32^ (Fig. 5d). Notably, DUX4 motifs preferred A in position 2, while DUXA-VP64, and sDUX preferred A or G equally in position 2 (Fig. 5d), in clear agreement with their respective luciferase reporter assays (Fig. 3b). Evaluating the simple pattern of two 4 nt cores of either TAAT or TGAT separated by any 3 nucleotides within peak sets revealed a clear preference for P3 TGAT by sDUX and DUXA-VP64, while P3 TAAT and N3 patterns were more frequent in DUX4 peaks, with the majority of DUX4 peaks bearing the N3 pattern (Fig. 5e). Enumerating the preferred motifs for each factor within their peak set revealed that the majority of dox-induced ATAC-seq peaks had at least one motif, with close to 50% having 2 or more (Fig. 5f).

Combining the RNA-seq with ATAC-seq to identify ATAC-seq peaks near genes upregulated by each factor, we evaluated the state of chromatin over each peak before and after dox treatment. Peaks that were induced only by DUX4 and not by sDUX or DUXA-VP64 tended to be completely closed prior to dox induction (Fig. 5g, second row). The same was not true for sDUX- or DUXA-VP64-specific peaks (Fig. 5g, first and third rows), or for shared peaks, which tended to be open prior to factor expression. This suggests that these factors are less able to bind regions of closed chromatin, while DUX4 has a potent pioneering activity. Interestingly, DUX4-specific peaks tended to have a weak signal for sDUX binding, suggesting that sDUX may also have a modicum of pioneer activity.

### DUXA inhibits DUX4 activity

Because *DUXA* is strongly induced by DUX4, because DUXA-VP64 acts on many of the same genes as DUX4, and because in its native form, DUXA lacks transcriptional activation potential, it seemed possible that DUXA might participate in negative feedback regulation of DUX4 activity by competing with DUX4 for binding sites that regulate key target genes. To directly investigate sites of DNA-binding of DUXA and DUX4, we performed ChIP-seq with epitope tagged versions of DUXA and DUX4 in 293T cells and evaluated peaks for overlap (Fig. 6a). As the ATAC-seq data and motif analysis suggested, there was substantial overlap in DNA-binding among these two factors (Fig. 6b, c). Approximately 70% of 1989 genes with a nearby DUX4 peak also showed a nearby DUXA peak (Fig. 6c).

**Figure 6.**
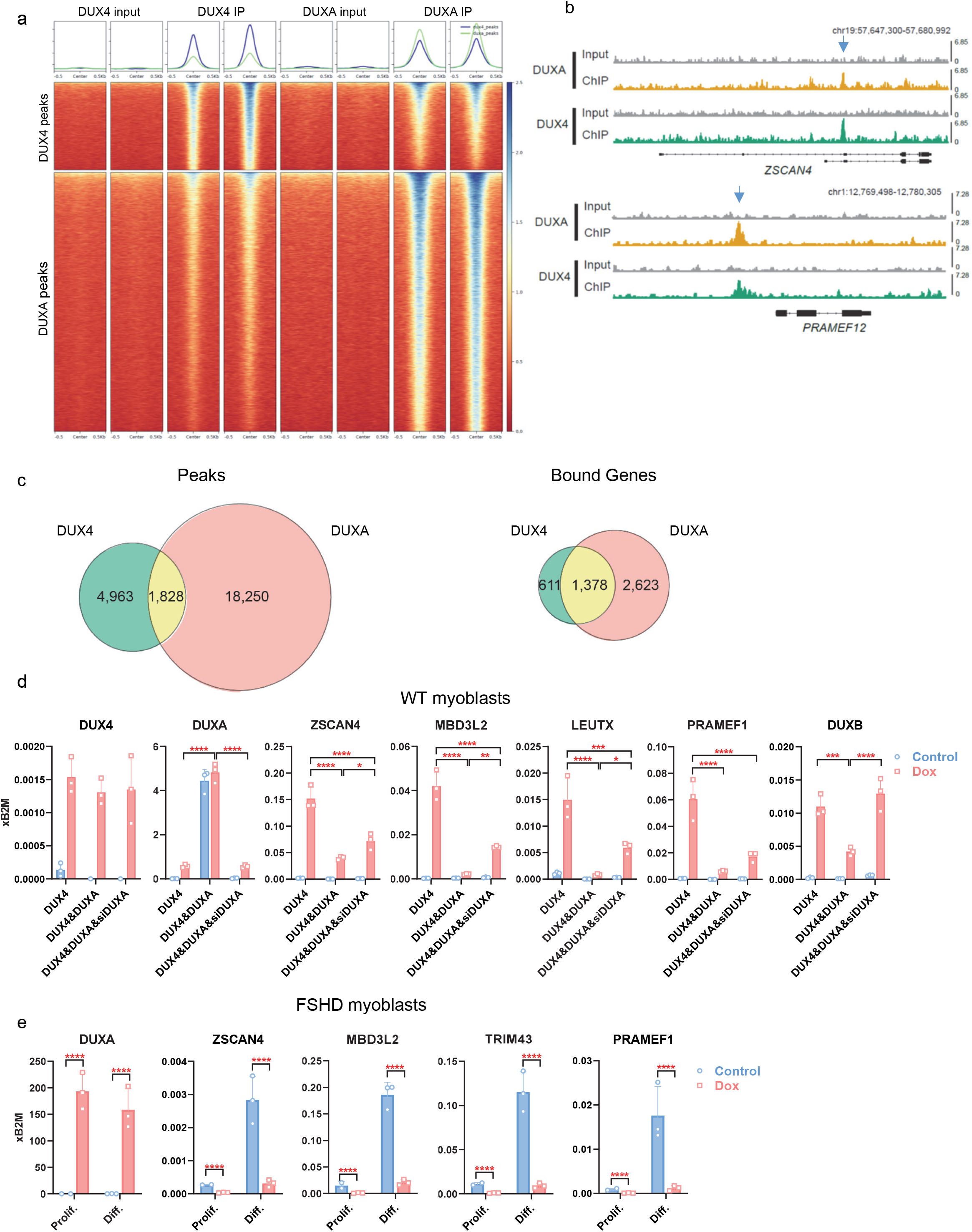
DUXA binds many DUX4 targets and inhibits DUX4 transcriptional activity. a. Tornado plots aligning on peak centers for called DUX4 peaks (upper panel) and DUXA peaks (lower panel) for both input and IP samples for both experiments. Overlap of DUX4 and DUXA peaks (peak calling: FDR <= 0.1, minimum treatment reads >= 1, enrichment >= 1, log10(qvalue) <= −0.5; not overlapping blacklisted regions like satellite repeats or misassemblies (ENCODE). b. ChIP-seq coverage tracks for DUXA and DUX4 surrounding the *ZSCAN4* (upper) and *PRAMEF12* (lower) genes. Input tracks are shown in gray and IP tracks are colored uniquely. Each track shown is a combination of two normalized biological replicates. The scale for each track is shown at the right and the gene structure is shown below. Arrows indicate shared peaks. c. Venn diagrams showing overlap in peaks (left) and in bound genes (right). Promoter = gene TSS + upstream 5000 nt + downstream 5000 nt; overlap of ≥ 1 nt. d. RTqPCR for DUX4 target genes in LHCN-M2iDUX4 immortalized human WT myoblasts co-expressing either DUXAiresEGFP or EGFP control, and cells in which overexpressed DUXA was knocked down by siRNA or control non targeted siRNA. Cells were induced with 200 ng/mL doxycycline for 8 hours. Data are presented as mean ± SEM; ***p<0.001, ****p<0.0001 by two-way ANOVA, n=3. e. RTqPCR for DUX4 target genes in immortalized human FSHD myoblasts engineered for dox-inducible for DUXA (M008-iDUXA). Cells were induced with 200 ng/mL doxycycline for 48 hours in proliferation conditions (Prolif.). For differentiation (Diff.), cells were cultured in differentiation medium for 3 days. DUXA was induced (200 ng/mL dox) over the last 24 hours of differentiation. Data are presented as mean ± SEM; ****p<0.0001 by two-way ANOVA, n=3.

We then tested whether DUXA could inhibit the activity of DUX4 by overexpressing DUXA constitutively in dox-inducible DUX4 expressing human myoblasts. Cells overexpressing DUXA showed a diminished ability of DUX4 to induce all of the targets we tested, compared to controls (Fig. 6d). As a further test of specificity, we knocked down DUXA in the constitutively expressing cell lines and observed the reversal of the DUXA effect. The activity of DUXA in suppressing DUX4 activity was independently replicated in another WT immortalized human myoblast cell line and in the context of pulsed DUX4 expression as opposed to continual expression (Supplementary Fig. 2).

### DUXA activity in FSHD patient-derived myoblasts

To evaluate the potential relevance of this effect in FSHD, we performed a similar experiment on an immortalized myoblast line from an FSHD-affected patient. In this case, the source of DUX4 is the endogenous DUX4 locus, and we delivered DUXA on a dox-inducible lentivirus. Because DUX4 target gene expression is strongly associated with differentiation, we differentiated these cells with doxycycline added and compared DUX4 target gene expression levels to those of cells differentiating under normal conditions (Fig. 6e). This revealed that also in FSHD cells, where DUX4 is expressed in a disease-relevant way, DUXA can inhibit the expression of DUX4 target genes.

## DISCUSSION

Because of the genetic divergence between the 3 clades of the DUX gene family, and the fact that the DUXC clade has acquired a tandem duplication amplification which has been retained (i.e., selected for) in all DUX family-bearing eutherian mammalian genomes evaluated ^2^, it seemed most probable that the double homeodomain proteins of the DUXC clade represent a neomorphic activity – a functional innovation over their ancestral single homeodomain progenitor. It was thus highly surprising that sDUX behaves like DUX4 in its cellular phenotypes of cytotoxicity and inhibition of myogenic differentiation and that, despite lacking C-terminal sequence conservation, sDUX has a C-terminal p300/CBP-dependent activation domain, and indeed has significantly overlapping downstream targets with DUX4, including a large number of ZGA-specific target genes. Rather than acquiring a neomorphic mutation, the DUXC clade seems to have retained the original function of sDUX, while the A and B clades represent loss of function or indeed antimorphic innovations.

The similarity in cytological phenotypes between sDUX and DUX4 is due in part to the similarity in the way these proteins bind DNA. Although sDUX has a single homeodomain, we find here that it binds DNA as a head-to-head dimer, in a nearly identical fashion to the head-to-head conformation of the tandem DUX4 homeodomains ^24^. This is in spite of a key difference, the lack of the interdomain linker of DUX4, which makes contacts with the DNA minor groove in the gap between the two core 4 nt sequences as well as at additional sites along the 11 nt motif (<u>TAAT</u>CTA<u>ATCA</u>), and which constrains the freedom of movement of the two HDs. Like DUX4, sDUX had no activity on HD core motifs spaced by 2 nt, suggesting that cooperative binding requires the 3 nt spacing. In this regard, it would be interesting to test whether DUX4 HD2 in isolation exhibits relaxed sequence selectivity as sDUX, and conversely, whether sDUX placed in a tandem homeodomain context would show higher selectivity than the non-covalent sDUX homodimer in DNA binding.

Regarding the specific sequence preferences of DUX proteins, it was previously demonstrated that although human and mouse DUXC proteins (DUX4 and DUX, resp.) recognize very similar 11 nt motifs, human DUX4 prefers the N3 (non-palindromic) TAAT---ATCA pattern against murine DUX, which prefers a P3 TGAT---ATCA pattern. The orientation of Glu70 in helix 3 of DUX4 HD1 specifies TAAT preference, and this orientation was variable in sDUX allowing it to bind TAAT and TGAT equally well. We infer that the relaxed sequence preference is an intrinsic feature of DUX proteins, likely derived from the ancestral HD, and that human and mouse DUXC specificity for the nonpalindromic *vs*. palindromic subsets is a derived feature. It will therefore be important to evaluate DUXC proteins from other species to determine whether evolved selectivity of motif recognition is an intrinsic feature of the DUXC clade, or whether human and mouse DUXC represent exceptions to the general pattern exemplified by sDUX. Irrespective of their idiosyncrasies, it is important to point out that both DUX4 and mouse DUX have the same basic cytotoxic phenotypes ^32,33^ and effects on differentiation ^33^, as indeed do sDUX and DUXA-VP64, whose sequence preferences encompass both those of DUX4 and DUX; thus the key function of these proteins is probably retained in their shared downstream targets, of which there are many ^14^, rather than those targets that are differentially regulated due to their evolved specificity.

If the DUXC clade retains the key properties of sDUX, what is the role of the DUXA and DUXB genes? Both DUXA and DUXB are transcriptionally inactive, however adding the VP64 activation domain revealed that DUXA acquired the DUX4 properties of cytotoxicity and inhibition of myogenic differentiation, and that it could regulate many DUX4 target genes, an interpretation strongly supported by the large overlap of DUXA and DUX4 ChIP-seq sites near DUX4-regulated genes. DUXB on the other hand was either unable to bind DNA in the context of our studies, or was not converted into an activator by fusion to VP64, as it showed extremely limited activity in these assays. Thus, despite DUX4, sDUX, and DUXA-VP64 each having many unique targets, they all 3 share a large set of common targets, suggesting that the common phenotype of cell death and inhibition of myogenic differentiation resides in this set of common transcriptional targets. DUXA in its native form of course fails to induce these targets. By binding a set of sites associated both with ZGA and with phenotypic cytotoxicity, but failing to induce them, DUXA inhibits the activity of DUX4. Importantly, DUXA is one of the most potently and rapidly upregulated targets of DUX4, suggesting a feedback-inhibitory mechanism. Such feedback inhibition would serve to limit the temporal effect of a burst of DUX4 expression, explaining why the burst of DUX4 target genes in ZGA is so brief. We note that such a negative feedback role is also demonstrated for mouse DUXBL against mouse DUX in a companion study ^34^. This feedback inhibition model stands in stark contrast to the previously suggested idea that DUXA serves to amplify or maintain the DUX4 signal after a burst of DUX4 ^23^. Although a positive feedback role for DUXA was attractive in explaining persistent effects of DUX4 long after a burst of expression, it would be incompatible with early embryogenesis and it is inconsistent with the data shown in the current study.

Inhibition of DUX4 by DUXA has implications for disease states involving DUX4. In FSHD, where DUX4 is potentially driving cytotoxicity or impairing myogenic regeneration, DUXA levels would serve as a rheostat on DUX4 activity and therefore limit the phenotype. Variation in DUXA levels among individuals, as well as among tissues, is thus of relevance in this disease of extremely variable penetrance and expressivity. Likewise, in cancers that show evidence of DUX4 expression ^20^, DUXA will serve to limit DUX4 activity on certain targets. The antagonistic relationship between DUXC and DUXA/DUXBL means that the mechanisms operating downstream of DUX4 in cancers in which DUX4 expression has been detected are not the same as the mechanisms that allow DUXBL to promote rhabdomyosarcoma ^21^.

These studies thus demonstrate that far from being a set of genes of similar activity, the DUX family of genes encodes proteins of opposing function, namely the DUXC clade with potent p300/CBP-dependent transcriptional activity, and the DUXA and B clades, which lack transcriptional activation potential, and in the case of DUXA, serve to competitively constrain in a feedback inhibitory manner, the activity of DUXC proteins like DUX4. This understanding serves to explain the extreme temporal limitation of activity of DUX4 and mouse DUX at ZGA and informs the interpretation of DUX4 and DUXA expression in cancer and FSHD.

## LIMITATIONS OF THE STUDY

This study investigates the activity of the human DUX proteins in cell lines cultured *in vitro*. As such, speculations about the function of these proteins in vivo are extrapolations. Formal proof of activity in humans would require *in vivo* experiments, ethically not feasible, although we do show that DUXA inhibits DUX4 in myoblasts from an FSHD patient (Fig. 6e). *In vivo* activity studies in another eutherian animal model such as mouse would provide valuable information, although outside the scope of the current study, which focuses on the human genes.

## MATERIALS AND METHODS

### Synthesis of expression constructs

The full length coding sequence for sDUX was identified by running the TBLASTN algorithm with the single platypus homeodomain sequence identified by Leidenroth and Hewitt against the platypus (mOrnAna1.p.v1) genome. The top hit (E-value = 5e-23) was found in the gene ENSOANG00000047618 located at chr3:113694167-113694310 adjacent to TMEM254 (CJ057) and ANXA11. A codon optimized version of the ENSOANG00000047618 open reading frame was synthesized by Biomatik. cDNAs for DUXA and DUXB were synthesized by Integrated DNA Technologies. FLAG and VP64 sequences were added by PCR amplification, and fragments were cloned into the pSAM2-Puro lentiviral vector ^13^ by In-Fusion HD Cloning (Takara).

### Cell Culture

Control LHCN-M2 immortalized human myoblasts ^35^ and FSHD (M008) immortalized human myoblasts were cultured in F10 medium (HyClone) with 20% FBS (Peak Serum), 2-mercaptoethanol 1x (Gibco), 10^−9^ M dexamethasone (Sigma), 10 ng/mL bFGF (Peprotech), Glutamax (Gibco), and Penicillin/Streptomycin (P/S, Gibco) at 37°C and 5% CO_2_. Immortalized M008 cells were derived by transducing primary myoblasts obtained from CD56 sorting of mononuclear cells from a muscle biopsy of an individual with FSHD under a protocol approved by the University of Minnesota IRB, with pbabe-cyclinD1+CDK4R24C (gift of Christopher Counter, Addgene) ^36^ and pLV-hTERT-IRES-hygro (gift of Tobias Meyer, Addgene) ^37^. The M007iDUX4 and M008iDUXA cell lines were generated from M007 and M008 immortalized myoblasts, which were derived by transduction of primary myoblasts from control and FSHD donors respectively, with two lentivectors, FUGW-rtTA and pSAM2-DUXA as previously described ^13^. The LHCN-M2iDUX4&DUXA cell line was generated by transducing LHCN-M2iDUX4 cells with pSAM-DUXA-ires-GFP lentivirus. GFP positive cells were FACS sorted two days post infection using a BD FACSAria (BD Biosciences, Franklin Lakes, NJ).

Myogenic differentiation was performed on gelatin (0.1%) coated dishes. Confluent cultures were washed once with PBS without Ca^2+^/Mg^2+^, and fed with differentiation medium (DMEM/F12 (Corning Cellgro) with 1x insulin/transferrin/selenium (Gibco), 1x NEAA (Gibco), Glutamax and P/S.

### RNA interference

Cells were seeded into 24-well plates (1.5× 10^5^/well), and the following day 50 nM siRNA for DUXA (L-034920-02-0005) or non-targeted control (D-001810-01-05) (SMARTpool, Dharmacon) were transfected using Lipofectamine RNAiMAX (Invitrogen). RNA was isolated 48 hours post-transfection and the effect on cell viability was analyzed at 72 and 96 hours post-transfection.

### EdU incorporation

EdU incorporation was measured using a Click-iT Plus EdU Flow Cytometry Assay Kit (Invitrogen). Cells were fixed and permeabilized in suspension, then incubated in Click-iT Plus reaction cocktail containing Alexa Fluor 647 picolyl azide. Flow cytometry was performed on a BD FACS Aria (BD Biosciences).

### Annexin V/7-AAD staining

Cells were trypsinized and stained with Annexin V and 7-AAD using APC Annexin V staining kit (BioLegend) according to the manufacturer’s instructions. Stained cells were evaluated on a FACSAria II (BD) and analyzed using FlowJo (FlowJo, LLC).

### Transduction

Viral supernatants were produced in 293T cells by transfection of lentivector DNA together with pVSVG and Δ8.9 packaging constructs, using Mirus TransIT-LTI transfection reagent (Mirus Bio). Medium was changed after 24 hours, and the viral supernatants were collected 48 hours post-transfection. Supernatants were then syringe-filtered (0.45 μm) and supplemented with 10 μg/mL polybrene (Millipore Sigma). Cells were incubated in viral supernatant diluted 1:2 overnight at 37°C, after which the supernatant was replaced with fresh culture medium. Infected cells were selected by Puromycin treatment (2.0 μg/mL) for 3 days.

### Cell viability (ATP) assay

Cells were plated in a 96 well dish (1 × 10^5^ cells/well), and the following day were induced with doxycycline. ATP assays were performed using CellTiter-Glo Luminescent Cell Viability Assay (Promega) according to the manufacturer’s instructions. Luminescence was analyzed on POLARstar Optima Microplate Reader (BMG Labtech, Offenburg, Germany).

### mRNA generation and transfection

Synthetic mRNAs were generated using mMESSAGE mMACHINE T7 ULTRA Transcription Kit (Invitrogen) using PCR product templates, which were produced by incorporating a T7 promoter sequence into the forward primers. Following ARCA-capped transcription and poly(A) tailing, synthetic RNA was purified and recovered using a QuickRNA miniprep kit (Zymo Research). Transfection of cells with mRNAs was achieved using a TransIT-mRNA transfection kit (Mirus Bio) according to the manufacturer’s instructions.

### Western blots

Cells were lysed with RIPA buffer supplemented with protease inhibitor cocktail (Complete, Roche), and proteins were separated on 10% SDS-PAGE gels, then transferred to PVDF membranes. Antibodies were diluted in 5% skim milk in TBST and incubated overnight at 4°C or 1 hour at RT. An appropriate HRP conjugated secondary antibody was incubated for 1 hour at RT. Membranes were then washed with TBST, and signal was visualized using Pierce ECL western blotting substrate (Thermo Scientific). Antibodies used in the study: GAPDH-HRP (1:5000, 60004, Proteintech), rabbit anti-Histone H3K18Ac (1:500, ab1191, Abcam), rabbit anti-Histone H3K27Ac (1:500, ab1791, Abcam, lot: GR3297878-1), anti-FLAG M2-Peroxidase (HRP, 1:10000, A8592, Sigma), and HRP conjugated anti-rabbit (1:5000, 111-035-003, Jackson Immuno Research, lot: 149393).

### RNA isolation, Quantitative Real Time RT-PCR (RTqPCR) and RNAseq

RNA was extracted using the Zymo RNA extraction kit following the manufacture’s protocol. cDNA was made using 0.5 μg total RNA with oligo-dT primer and Verso cDNA Synthesis Kit (Thermo Scientific) following the manufacturer’s instructions. qPCR was performed by using Premix Probe Ex Taq or SYBR-green (Takara) and commercially available probes from Applied Biosystems (*GAPDH*, Hs99999905_m1; *B2M*, Hs00187842_m1; *ZSCAN*, Hs00357549_m1; *LEUTX*, Hs01028718_m1; *PRAMEF1*, Hs04401269, *TRIM43*, Hs00299174_m1, *MYH7*, Hs01110602_m1, *DUX4*, Hs03037970_g1, *MBD3L2*, Hs00544743_m1) or custom-designed primers (*DUXA*: F: 5’ TCT TGC CCT GCT CTT CTT GT and R: 5’ CCT GGG ATT GAT TCC AGA GA; *DUXB* F: 5’ CCC TGA TAA AGC TGC CAG AG and R: 5’ TGA GTC AGA TGC TGG GAC TG; *RFPL4B* F: 5’ GGC TGA ATT CAA GTG GGT CT and R: 5’ GAG ACG TAG GCT TCG GAT CTT). Gene expression levels were normalized to that of *GAPDH* or *B2M* and analyzed with 7500 System Software using the ΔCT method (Applied Biosystems).

RNA-seq library preparation was done with 500 ng total RNA. One set of libraries (Fig. 3) was prepared using the Swift Rapid RNA Library Kit (SwiftBioscience) and 36 base paired-end sequenced on an Illumina NextSeq at the University of Minnesota Genomics Center (UMGC). The other set (Fig. 4) was prepared and sequenced (2 × 150 paired-end) on an Illumina HiSeq at Genewiz (NJ, USA). Conventional gene expression was quantified using Salmon (v1.9) with the human Gencode v38 transcriptome modified to contain the sDUX coding sequence. Analysis of expression of repetitive elements was performed by mapping trimmed reads to the GRCh38 genome with Star (v2.7.2) and enumerating the reads mapping to UCSC RMSK features using Rsubread allowing for fractional counts. Gene expression differences for conventional genes and repetitive elements was performed with DESeq2 (v1.36.0).

### ATAC-seq

For each construct, biological duplicates from doxycycline-treated and untreated myoblasts were collected using Trypsin-EDTA followed by dilution with culture medium and centrifugation at 400g for 5 minutes. 50,000 cells were washed with 200 μL of cold PBS then resuspended in 100 μL of cold lysis buffer (10mM Tris-HCl pH 7.4, 10mM NaCl, 3mM MgCl2, 0.1% IGEPAL CA-630), spun at 500 g for 10 minutes at 4°C and resuspended in 50 μL of the transposition reaction mix. Transposition occurred at 37°C for 30 minutes, after which transposed DNA was purified using a Qiagen MinElute Kit and eluted in 10μL Elution Buffer. Chromatin accessibility studies were performed following the protocol described by ^38^. PCR amplification using Illumina-compatible adapter-barcodes and final library preparation were performed at the UMGC. After Quality Control, libraries were pooled and sequenced on NovaSeq S4 2×150-bp run (Illumina). Mapping to the human hg38 genome and peak calling were performed using the ENCODE ATAC-seq pipeline.

### ChIP-seq sample and library preparation

HEK293 cells were transfected with Doxycycline inducible plasmids either encoding for V5 epitope-tagged DUX4 or DUXA. 24 h after later, gene expression was induced by addition of Doxycycline (Sigma, D9891-10G) to a final concentration of 100 ng/ml for 24 h. Next, chromatin-protein complexes were cross-linked by addition of methanol-free formaldehyde (ThermoFisher, 28906) at a final concentration of 1 % for 10 min at RT. Cells were then washed three times in PBS and chromatin was isolated and sheared to 200-700 bp using the truChIP Chromatin Shearing Kit (Covaris, 520154) and a Covaris E220evolution focused-ultrasonicator following the manufacturer’s recommendations. Subsequently, 30 μg of chromatin was diluted 1:4 with RIPA-buffer (10 mM Tris-HCl pH 8.0, 1 mM EDTA pH 8.0, 140 mM NaCl, 1 % Triton X-100, 0.1 % sodium deoxycholate) and incubated with 4 μg anti-V5-antibody (Abcam, ab9116) at 4° C overnight. Antibody-bound chromatin was then coupled to protein A beads (Diagenode, C03020002) for three hours at 4° C. Samples were then washed four times with RIPA-buffer, two times with high-salt-RIPA-buffer (10 mM Tris-HCl pH 8.0, 1 mM EDTA pH 8.0, 500 mM NaCl, 1 % Triton X-100, 0.1 % sodium deoxycholate), two times with LiCl-buffer (10 mM Tris-HCl pH 8.0, 1 mM EDTA pH 8.0, 250 mM LiCl, 0.5 % NP-40, 0.5 % sodium deoxycholate) and finally two times with TE Buffer (10 mM Tris-HCl pH 8.0, 1 mM EDTA pH 8.0). RNA was eliminated by addition of RNAseA (Thermo Fisher Scientific, EN0531) to a final concentration of 0.1 mg/ml in 10 mM Tris-HCl pH 8.0, 1 mM EDTA pH 8.0, 300 mM NaCl, 0.5 % SDS and incubated at 37° C for 30 min. Reverse crosslinking was achieved by addition of Proteinase K (ThermoFisher, EO0491) to a final concentration of 0.4 mg/ml and incubation at 55° C for 1 h and then at 65° C overnight. DNA was purified using the NucleoSpin Gel and PCR Clean-up kit (Macherey-Nagel, 740609) and quantified by Qubit dsDNA HS Assay Kit (Thermo Fisher Scientific). 2-3 ng of DNA was used as input for TruSeq ChIP Library Preparation Kit (Illumina) with following modifications. Instead of gel-based size selection before final PCR step, libraries were size selected by SPRI-bead based approach after final PCR with 18 cycles. In detail, samples were 1^st^ cleaned up by 1x bead:DNA ratio to eliminate residuals from PCR reaction, followed by 2-sided-bead cleanup step with initially 0.6x bead:DNA ratio to exclude larger fragments. Supernatant was transferred to new tube and incubated with additional beads in 0.2x bead:DNA ratio for eliminating smaller fragments, like adapter and primer dimers. Bound DNA samples were washed with 80% ethanol, dried and resupended in TE buffer. Library integrity was verified with LabChip Gx Touch 24 (Perkin Elmer). Sequencing was performed on the NextSeq500 instrument (Illumina) using v2 chemistry with 1×75bp single end setup.

### ChIP-seq analysis

Trimmomatic version 0.39 was employed to trim reads after a quality drop below a mean of Q20 in a window of 20 nucleotides and keeping only filtered reads longer than 15 nucleotides (Bolger et al., Trimmomatic: a flexible trimmer for Illumina sequence data). Reads were aligned versus Ensembl human genome version hg38 (Ensembl release 104) with STAR 2.7.10a (Dobin et al., STAR: ultrafast universal RNA-seq aligner). Aligned reads were filtered to remove: duplicates with Picard 2.27.1 (Picard: A set of tools (in Java) for working with next generation sequencing data in the BAM format), spliced, multi-mapping, ribosomal, or mitochondrial reads. Peak calling was performed with Macs version 3.0.0a6 with FDR < 0.1 and “--scale-to large” (Zhang et al., Model-based Analysis of ChIP-Seq). Peaks overlapping ENCODE blacklisted regions (known misassemblies, satellite repeats) were excluded. Remaining peaks were unified to represent a common set of regions for all samples and counts were produced with bigWigAverageOverBed (UCSC Toolkit). The raw count matrix was normalized with DESeq2 version 1.30.1 (Love et al., Moderated estimation of fold change and dispersion for RNA-Seq data with DESeq2). Peaks were annotated with the gene + promoter (TSS +-5000 nt) with the largest overlap based on Ensembl release 104. Contrasts were created with DESeq2 based on the normalized union peak matrix with size factors set to zero. Peaks were classified as significantly differential at average count > 10 and −1 < log2FC > 1.

### Immunofluorescence

Cells were fixed in 4% PFA for 10 min., washed twice with PBS, permeabilized with 0.3% Triton X for 30 min, and blocked with 3% BSA for 1 hour at RT. Primary antibody (MF20, 1:20, Hybridoma bank, University of Iowa) was diluted in 3% BSA and incubated ON at 4 °C. Secondary antibody (Alexa fluor 555 Goat Anti-Mouse, 1:500, Invitrogen) was applied for 60 min at RT. Nuclei were visualized using DAPI (1:5000, Sigma).

### Biochemical and structural studies

The homeodomain of platypus sDUX was expressed as MBP-fusion from a codon-optimized gene cloned into the pMAL-c5x bacterial expression vector. The construct contained an 8xHis-tag and the Human Rhinovirus (HRV) 3C Protease cleavage site between MBP and sDUX. The protein was expressed in the *E. coli* strain BL21(DE3) and purified from the soluble bacterial extract using Ni-NTA affinity chromatography. After a treatment with HRV 3C protease, sDUX was further purified over a Superdex 75 size-exclusion column operating with 20 mM Tris-HCl, pH 7.4, 0.5 M NaCl, 5 mM β-mercaptoethanol, concentrated by ultrafiltration, flash-frozen in liquid nitrogen and stored at –80 °C. The protein concentration was determined based on UV absorbance. The amino acid sequence of the purified sDUX HD was: GPAREGARRK RTTFNKTQLE ILVKSFNKDP YPGIGVREHL ASLIQIPESR IQVWFQNRRA RQLGQKKKLEV. DUX4 HD1-HD2 was prepared and EMSA conducted as described previously ^24^.

For crystallographic studies, sDUX-DNA complex was prepared by mixing the purified protein with a 17 bp dsDNA substrate (5′-GCG TAA TCT AAT CAA CA-3′ / 5′-TGT TGA TTA GAT TAC GC-3′) at a molar ratio of 2:1 in 10 mM Tris-HCl, pH 7.4, 0.1 M NaCl, at a protein concentration of 20 mg ml^-1^. The complex was crystallized in sitting drop vapor diffusion mode using the reservoir solution consisting of 0.2 M calcium chloride, 50 mM HEPES buffer, pH 7.5, 28% polyethylene glycol 400, 2 mM spermine. The sDUX-DNA crystals were cryo-protected by brief soaking in the reservoir solution supplemented with 20 % ethylene glycol and flash cooled by plunging in liquid nitrogen. X-ray diffraction data were collected at the NE-CAT beamline 24-ID-C of the Advanced Photon Source (Lemont, IL) and processed using XDS ^39^. The structures were determined by molecular replacement with PHASER^40^ using the DUX4-DNA complex (PDB ID: 6E8C) ^24^ as the search model. Iterative model building and refinement were conducted using COOT ^41^ and PHENIX ^42^. To determine the identity of DNA strands in the pseudo-symmetrical complex, we collected additional data set on a crystal grown with the bottom strand containing a 5-bromouracil substitution (5′-TG/i5Br-dU/ TGA TTA GAT TAC GC-3′), which gave an improved resolution albeit with strong anisotropy. A summary of crystallographic data statistics is shown in Supplementary Table 1. Figures were generated using PyMOL (https://pymol.org/2/).

### Statistics

All experiments were repeated in at least in three biological replicates. Significance were calculated by one- or two-way ANOVA with GraphPad; **** indicates p<0.0001, *** p<0.001, ** p<0.01, and * p<0.05.

## ACKNOWLEDGEMENTS

We thank Jasmine Gulik, Ana Mitanoska, David Oyler, Natalie Xu and MacKenzie Molina for the help with molecular analyses. This work was supported by a grant from the National Institute of Arthritis and Musculoskeletal and Skin Diseases (R01 AR055685 to M.K.) and the National Institute of General Medical Sciences (R35 GM118047 to H.A.). X-ray diffraction data were collected at the Northeastern Collaborative Access Team beamlines, which are funded by the US National Institutes of Health (NIGMS P30 GM124165). The Pilatus 6M detector on 24-ID-C beamline is funded by a NIH-ORIP HEI grant (S10 RR029205). This research used resources of the Advanced Photon Source, a U.S. Department of Energy (DOE) Office of Science User Facility operated for the DOE Office of Science by Argonne National Laboratory under Contract No. DE-AC02-06CH11357. The monoclonal antibody against MHC was obtained from the Developmental Studies Hybridoma Bank, developed under the auspices of the NICHD and maintained by the University of Iowa.

## Author Contributions

Experimentation: DB, EAT, ETE, MDG, LY, FL, AM, KS Writing: DB, MDG, JK, HA, MK

## Competing financial and other interests

The authors have no competing financial or other interests to declare. The authors have declared that no conflict of interest exists.

## Ethics Statement

Human subjects work was performed in accordance with a protocol approved by the University of Minnesota IRB.

## Data Availability

Atomic coordinates and structure factors have been deposited in the Protein Data Bank (PDB) under accession codes 8EJO and 8EJP. RNA-seq, ATAC-seq, and ChIP-seq data have been submitted to GEO (GSE214245 and GSE214230).

## EXTENDED DATA LEGENDS

**Supplementary Figure 1. Controls (uninduced cells) for luciferase assays**

Luciferase assays in uninduced 293T cells with various dox-inducible DUX family proteins and VP64 derivatives on the 3 flavors of DUX motif (N3 and P3) and the Pax7 (P2-type) motifs.

**Supplementary Figure 2. DUXA suppress DUX4 transcriptional activity**

a. Pulse induction in WT myoblasts. RTqPCR for DUX4 target genes in LHCNiDUX4 immortalized human WT myoblasts co-expressing either DUXAiresEGFP or EGFP control, and cells in which overexpressed DUXA was knocked down by siRNA or control non targeted siRNA. Cells were pulse-induced with 200 ng/ml doxycycline for 1 hours and analyzed 8 hours later. Data are presented as mean ± SEM; **p<0.01, ****p<0.0001 by two-way ANOVA, n=3.

b. RTqPCR for DUX4 target genes in WT human immortalized M007iDUX4 myoblasts. Cells were transfected with *DUXA* or *GFP* mRNA for 24 hours, and over the last 4 hours, cells were induced with 200 ng/mL doxycycline. Data are presented as mean ± SEM; *p<0.05, **p<0.01, ****p<0.0001 by two-way ANOVA, n=3.

## DESCRIPTION OF SUPPLEMENTARY FILES

***Supplementary Figure 1***

Luciferase Assay Controls

***Supplementary Figure 2***

DUXA competition in DUX4-induced M007 and DUX4-pulsed LHCN-M2 immortalized human myoblasts.

***Supplementary Table 1***

Crystallographic data statistics

## Notes

### Competing Interest Statement

The authors have declared no competing interest.

## REFERENCES

1. Clapp, J., Mitchell, L.M., Bolland, D.J., Fantes, J., Corcoran, A.E., Scotting, P.J., Armour, J.A., and Hewitt, J.E. (2007). Evolutionary conservation of a coding function for D4Z4, the tandem DNA repeat mutated in facioscapulohumeral muscular dystrophy. Am J Hum Genet 81, 264–279.

2. Leidenroth, A., and Hewitt, J.E. (2010). A family history of DUX4: phylogenetic analysis of DUXA, B, C and Duxbl reveals the ancestral DUX gene. BMC Evol Biol 10, 364. 1471-2148-10-364 [pii] 10.1186/1471-2148-10-364.

3. Gabriels, J., Beckers, M.C., Ding, H., De Vriese, A., Plaisance, S., van der Maarel, S.M., Padberg, G.W., Frants, R.R., Hewitt, J.E., Collen, D., and Belayew, A. (1999). Nucleotide sequence of the partially deleted D4Z4 locus in a patient with FSHD identifies a putative gene within each 3.3 kb element. Gene 236, 25–32. S0378-1119(99)00267-X [pii].

4. van Overveld, P.G., Lemmers, R.J., Sandkuijl, L.A., Enthoven, L., Winokur, S.T., Bakels, F., Padberg, G.W., van Ommen, G.J., Frants, R.R., and van der Maarel, S.M. (2003). Hypomethylation of D4Z4 in 4q-linked and non-4q-linked facioscapulohumeral muscular dystrophy. Nature genetics 35, 315–317.

5. Snider, L., Geng, L.N., Lemmers, R.J., Kyba, M., Ware, C.B., Nelson, A.M., Tawil, R., Filippova, G.N., van der Maarel, S.M., Tapscott, S.J., and Miller, D.G. (2010). Facioscapulohumeral dystrophy: incomplete suppression of a retrotransposed gene. PLoS Genet 6, e1001181. 10.1371/journal.pgen.1001181.

6. Wijmenga, C., Hewitt, J.E., Sandkuijl, L.A., Clark, L.N., Wright, T.J., Dauwerse, H.G., Gruter, A.M., Hofker, M.H., Moerer, P., Williamson, R., and J., v.O.G. (1992). Chromosome 4q DNA rearrangements associated with facioscapulohumeral muscular dystrophy. Nature genetics 2, 26–30.

7. Kowaljow, V., Marcowycz, A., Ansseau, E., Conde, C.B., Sauvage, S., Matteotti, C., Arias, C., Corona, E.D., Nunez, N.G., Leo, O., et al. (2007). The DUX4 gene at the FSHD1A locus encodes a pro-apoptotic protein. Neuromuscular disorders : NMD 17, 611–623.

8. Bosnakovski, D., Xu, Z., Gang, E.J., Galindo, C.L., Liu, M., Simsek, T., Garner, H.R., Agha-Mohammadi, S., Tassin, A., Coppee, F., et al. (2008). An isogenetic myoblast expression screen identifies DUX4-mediated FSHD-associated molecular pathologies. EMBO J 27, 2766–2779. emboj2008201 [pii] 10.1038/emboj.2008.201.

9. Bosnakovski, D., Gearhart, M.D., Toso, E.A., Ener, E.T., Choi, S.H., and Kyba, M. (2018). Low level DUX4 expression disrupts myogenesis through deregulation of myogenic gene expression. Scientific reports 8, 16957. 10.1038/s41598-018-35150-8.

10. Geng, L.N., Yao, Z., Snider, L., Fong, A.P., Cech, J.N., Young, J.M., van der Maarel, S.M., Ruzzo, W.L., Gentleman, R.C., Tawil, R., and Tapscott, S.J. (2012). DUX4 activates germline genes, retroelements, and immune mediators: implications for facioscapulohumeral dystrophy. Dev Cell 22, 38–51. S1534-5807(11)00523-5 [pii] 10.1016/j.devcel.2011.11.013.

11. Kawamura-Saito, M., Yamazaki, Y., Kaneko, K., Kawaguchi, N., Kanda, H., Mukai, H., Gotoh, T., Motoi, T., Fukayama, M., Aburatani, H., et al. (2006). Fusion between CIC and DUX4 up-regulates PEA3 family genes in Ewing-like sarcomas with t(4;19)(q35;q13) translocation. Hum Mol Genet 15, 2125–2137. 10.1093/hmg/ddl136.

12. Bosnakovski, D., Lamb, S., Simsek, T., Xu, Z., Belayew, A., Perlingeiro, R., and Kyba, M. (2008). DUX4c, an FSHD candidate gene, interferes with myogenic regulators and abolishes myoblast differentiation. Exp Neurol 214, 87–96. S0014-4886(08)00297-5 [pii] 10.1016/j.expneurol.2008.07.022.

13. Choi, S.H., Gearhart, M.D., Cui, Z., Bosnakovski, D., Kim, M., Schennum, N., and Kyba, M. (2016). DUX4 recruits p300/CBP through its C-terminus and induces global H3K27 acetylation changes. Nucleic Acids Res 44, 5161–5173. 10.1093/nar/gkw141.

14. Hendrickson, P.G., Dorais, J.A., Grow, E.J., Whiddon, J.L., Lim, J.W., Wike, C.L., Weaver, B.D., Pflueger, C., Emery, B.R., Wilcox, A.L., et al. (2017). Conserved roles of mouse DUX and human DUX4 in activating cleavage-stage genes and MERVL/HERVL retrotransposons. Nature genetics 49, 925–934. 10.1038/ng.3844.

15. Whiddon, J.L., Langford, A.T., Wong, C.J., Zhong, J.W., and Tapscott, S.J. (2017). Conservation and innovation in the DUX4-family gene network. Nature genetics 49, 935–940. 10.1038/ng.3846.

16. De Iaco, A., Planet, E., Coluccio, A., Verp, S., Duc, J., and Trono, D. (2017). DUX-family transcription factors regulate zygotic genome activation in placental mammals. Nature genetics 49, 941–945. 10.1038/ng.3858.

17. Chen, Z., and Zhang, Y. (2019). Loss of DUX causes minor defects in zygotic genome activation and is compatible with mouse development. Nature genetics 51, 947–951. 10.1038/s41588-019-0418-7.

18. Bosnakovski, D., Gearhart, M.D., Ho Choi, S., and Kyba, M. (2021). Dux facilitates post-implantation development, but is not essential for zygotic genome activation. Biol Reprod 104, 83–93. 10.1093/biolre/ioaa179.

19. De Iaco, A., Verp, S., Offner, S., Grun, D., and Trono, D. (2020). DUX is a non-essential synchronizer of zygotic genome activation. Development 147, dev177725. 10.1242/dev.177725.

20. Chew, G.L., Campbell, A.E., De Neef, E., Sutliff, N.A., Shadle, S.C., Tapscott, S.J., and Bradley, R.K. (2019). DUX4 Suppresses MHC Class I to Promote Cancer Immune Evasion and Resistance to Checkpoint Blockade. Dev Cell 50, 658–671 e657. 10.1016/j.devcel.2019.06.011.

21. Preussner, J., Zhong, J., Sreenivasan, K., Gunther, S., Engleitner, T., Kunne, C., Glatzel, M., Rad, R., Looso, M., Braun, T., and Kim, J. (2018). Oncogenic Amplification of Zygotic Dux Factors in Regenerating p53-Deficient Muscle Stem Cells Defines a Molecular Cancer Subtype. Cell stem cell 23, 794–805 e794. 10.1016/j.stem.2018.10.011.

22. Bosnakovski, D., Oyler, D., Mitanoska, A., Douglas, M., Ener, E.T., Shams, A.S., and Kyba, M. (2022). Persistent Fibroadipogenic Progenitor Expansion Following Transient DUX4 Expression Provokes a Profibrotic State in a Mouse Model for FSHD. Int J Mol Sci 23. 10.3390/ijms23041983.

23. Jiang, S., Williams, K., Kong, X., Zeng, W., Nguyen, N.V., Ma, X., Tawil, R., Yokomori, K., and Mortazavi, A. (2020). Single-nucleus RNA-seq identifies divergent populations of FSHD2 myotube nuclei. PLoS Genet 16, e1008754. 10.1371/journal.pgen.1008754.

24. Lee, J.K., Bosnakovski, D., Toso, E.A., Dinh, T., Banerjee, S., Bohl, T.E., Shi, K., Orellana, K., Kyba, M., and Aihara, H. (2018). Crystal Structure of the Double Homeodomain of DUX4 in Complex with DNA. Cell reports 25, 2955–2962 e2953. 10.1016/j.celrep.2018.11.060.

25. Banerji, C.R.S., Panamarova, M., Hebaishi, H., White, R.B., Relaix, F., Severini, S., and Zammit, P.S. (2017). PAX7 target genes are globally repressed in facioscapulohumeral muscular dystrophy skeletal muscle. Nat Commun 8, 2152. 10.1038/s41467-017-01200-4.

26. Banerji, C.R.S., and Zammit, P.S. (2019). PAX7 target gene repression is a superior FSHD biomarker than DUX4 target gene activation, associating with pathological severity and identifying FSHD at the single-cell level. Hum Mol Genet 28, 2224–2236. 10.1093/hmg/ddz043.

27. Bosnakovski, D., da Silva, M.T., Sunny, S.T., Ener, E.T., Toso, E.A., Yuan, C., Cui, Z., Walters, M.A., Jadhav, A., and Kyba, M. (2019). A novel P300 inhibitor reverses DUX4-mediated global histone H3 hyperacetylation, target gene expression, and cell death. Sci Adv 5, eaaw7781. 10.1126/sciadv.aaw7781.

28. Bosnakovski, D., Ener, E.T., Cooper, M.S., Gearhart, M.D., Knights, K.A., Xu, N.C., Palumbo, C.A., Toso, E.A., Marsh, G.P., Maple, H.J., and Kyba, M. (2021). Inactivation of the CIC-DUX4 oncogene through P300/CBP inhibition, a therapeutic approach for CIC-DUX4 sarcoma. Oncogenesis 10, 68. 10.1038/s41389-021-00357-4.

29. Beerli, R.R., Segal, D.J., Dreier, B., and Barbas, C.F., 3rd (1998). Toward controlling gene expression at will: specific regulation of the erbB-2/HER-2 promoter by using polydactyl zinc finger proteins constructed from modular building blocks. Proceedings of the National Academy of Sciences of the United States of America 95, 14628–14633. 10.1073/pnas.95.25.14628.

30. Yu, X., Liang, S., Chen, M., Yu, H., Li, R., Qu, Y., Kong, X., Guo, R., Zheng, R., Izsvak, Z., et al. (2022). Recapitulating early human development with 8C-like cells. Cell reports 39, 110994. 10.1016/j.celrep.2022.110994.

31. Peaston, A.E., Evsikov, A.V., Graber, J.H., de Vries, W.N., Holbrook, A.E., Solter, D., and Knowles, B.B. (2004). Retrotransposons regulate host genes in mouse oocytes and preimplantation embryos. Dev Cell 7, 597–606. 10.1016/j.devcel.2004.09.004.

32. Eidahl, J.O., Giesige, C.R., Domire, J.S., Wallace, L.M., Fowler, A.M., Guckes, S.M., Garwick-Coppens, S.E., Labhart, P., and Harper, S.Q. (2016). Mouse Dux is myotoxic and shares partial functional homology with its human paralog DUX4. Hum Mol Genet 25, 4577–4589. 10.1093/hmg/ddw287.

33. Bosnakovski, D., Daughters, R.S., Xu, Z., Slack, J.M., and Kyba, M. (2009). Biphasic myopathic phenotype of mouse DUX, an ORF within conserved FSHD-related repeats. PLoS One 4, e7003. 10.1371/journal.pone.0007003.

34. Vega-Sendino, M., Olbrich, T., Stein, P., Tillo, D., Carey, G.I., Virginia, Savy, Saykali, B., Domingo, C.N., Maity, T.K., et al. (2022). The homeobox transcription factor DUXBL controls exit from totipotency. Preprint at BioRxiv https://www.biorxiv.org/content/10.1101/2022.09.19.508541v1.

35. Zhu, C.H., Mouly, V., Cooper, R.N., Mamchaoui, K., Bigot, A., Shay, J.W., Di Santo, J.P., Butler-Browne, G.S., and Wright, W.E. (2007). Cellular senescence in human myoblasts is overcome by human telomerase reverse transcriptase and cyclin-dependent kinase 4: consequences in aging muscle and therapeutic strategies for muscular dystrophies. Aging Cell 6, 515–523. 10.1111/j.1474-9726.2007.00306.x.

36. Kendall, S.D., Linardic, C.M., Adam, S.J., and Counter, C.M. (2005). A network of genetic events sufficient to convert normal human cells to a tumorigenic state. Cancer Res 65, 9824–9828. 10.1158/0008-5472.CAN-05-1543.

37. Hayer, A., Shao, L., Chung, M., Joubert, L.M., Yang, H.W., Tsai, F.C., Bisaria, A., Betzig, E., and Meyer, T. (2016). Engulfed cadherin fingers are polarized junctional structures between collectively migrating endothelial cells. Nat Cell Biol 18, 1311–1323. 10.1038/ncb3438.

38. Buenrostro, J.D., Wu, B., Chang, H.Y., and Greenleaf, W.J. (2015). ATAC-seq: A Method for Assaying Chromatin Accessibility Genome-Wide. Curr Protoc Mol Biol 109, 21 29 21–21 29 29. 10.1002/0471142727.mb2129s109.

39. Kabsch, W. (2010). Xds. Acta Crystallogr D Biol Crystallogr 66, 125–132. 10.1107/S0907444909047337.

40. McCoy, A.J., Grosse-Kunstleve, R.W., Adams, P.D., Winn, M.D., Storoni, L.C., and Read, R.J. (2007). Phaser crystallographic software. J Appl Crystallogr 40, 658–674. 10.1107/S0021889807021206.

41. Emsley, P., Lohkamp, B., Scott, W.G., and Cowtan, K. (2010). Features and development of Coot. Acta Crystallogr D Biol Crystallogr 66, 486–501. 10.1107/S0907444910007493.

42. Adams, P.D., Afonine, P.V., Bunkoczi, G., Chen, V.B., Davis, I.W., Echols, N., Headd, J.J., Hung, L.W., Kapral, G.J., Grosse-Kunstleve, R.W., et al. (2010). PHENIX: a comprehensive Python-based system for macromolecular structure solution. Acta Crystallogr D Biol Crystallogr 66, 213–221. 10.1107/S0907444909052925.

