## Supplementary table and figures for "Antagonism among DUX family members evolved from an ancestral toxic single homeodomain protein"

### Supplementary Figure 1

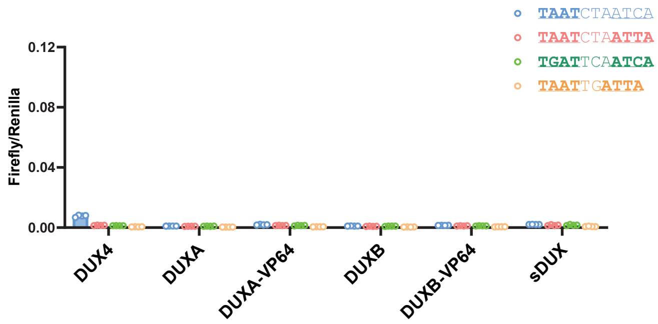


Luciferase assays in uninduced 293T cells with various dox-inducible DUX family proteins and VP64 derivatives on the 3 flavors of DUX motif (N3 and P3) and the Pax7 (P2-type) motifs.

Supplementary Figure 2

**a**

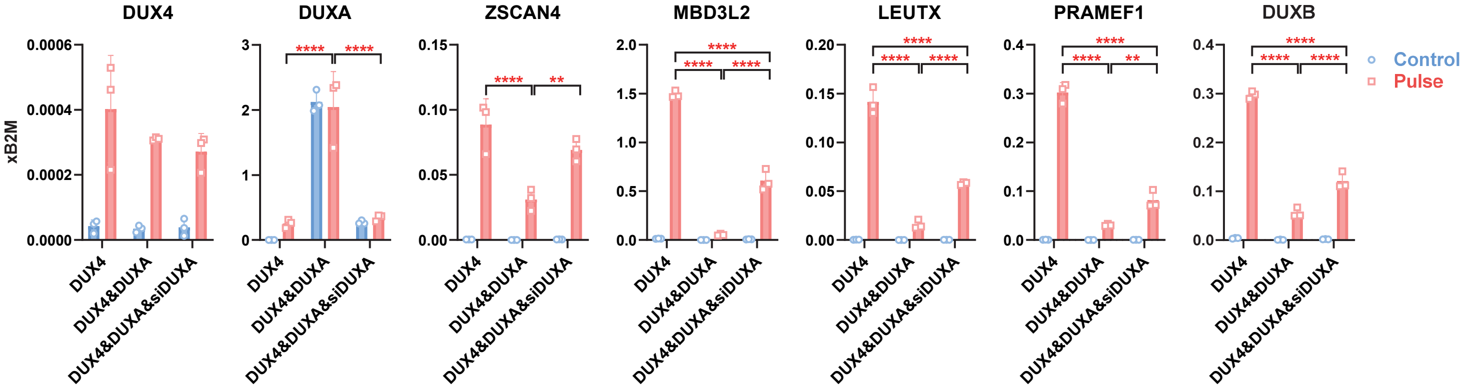


**b**

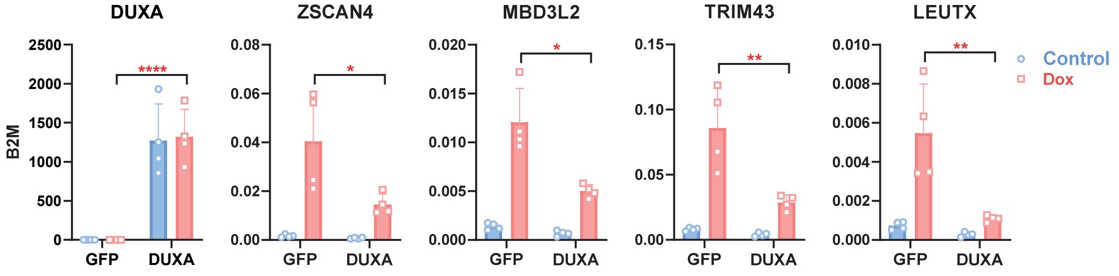


a. Pulse induction in WT myoblasts. RTqPCR for DUX4 target genes in LHCN-iDUX4 immortalized human WT myoblasts co-expressing either DUXAiresEGFP or EGFP control, and cells in which overexpressed DUXA was knocked down by siRNA or control non targeted siRNA. Cells were pulse-induced with 200 ng/ml doxycycline for 1 hours and analyzed 8 hours later. Data are presented as mean ± SEM; **$p<0.01$, ****$p<0.0001$ by two-way ANOVA, n=3..

b. RTqPCR for DUX4 target genes in WT human immortalized M007-iDUX4 myoblasts. Cells were transfected with DUXA or GFP mRNA for 24 hours, and over the last 4 hours, cells were induced with 200 ng/mL doxycycline. Data are presented as mean ± SEM; *$p<0.05$, **$p<0.01$, ****$p<0.0001$ by two-way ANOVA, n=3.

**Table 1. Data collection and refinement statistics.**

| | sDUX4/DNA (8EJO) | sDUX4/BrU-DNA (8EJP) |
|---|---|---|
| **Data collection** | | |
| Wavelength (Å) | 0.979 | 0.979 |
| Resolution range (Å) | 74.20 - 2.67 (2.80 - 2.67) | 59.68 - 2.17 (2.21 - 2.17)† |
| Space group | $P4_32_12$ | $P4_32_12$ |
| Unit cell | | |
| *a, b, c* (Å) | 74.02, 74.02, 97.31 | 74.50, 74.50, 99.69 |
| Total reflections | 50642 (6516) | 215341 (11037) |
| Unique reflections | 8096 (1021) | 15354 (747) |
| Multiplicity | 6.3 (6.4) | 14.0 (14.8) |
| Completeness (%) | 99.20 (98.00) | 99.90 (100.0) |
| $<I/\sigma(I)>$ | 14.30 (1.00) | a*,b* 7.40 (0.40)/c* 10.7 (1.5) |
| $R_{merge}$ | 0.059 (1.15) | 0.131 (6.20) |
| $R_{meas}$ | 0.064 (1.37) | 0.132 (6.47) |
| $R_{p.i.m.}$ | 0.032 (0.72) | 0.051 (2.33) |
| $CC_{1/2}$ | 0.999 (0.54) | a*,b* 0.998(0.121)/c* 0.998(0.649) |
| **Refinement** | | |
| Reflections for $R_{work}$ | 7147 (401) | 10466 (161) |
| Reflections for $R_{free}$ | 348 (15) | 545 (10) |
| $R_{work}/R_{free}$ | 0.206/0.247 | 0.227/0.259 |
| No. of non-H atoms | 1655 | 1658 |
| Macromolecules | 1648 | 1648 |
| Ligands | 4 | 8 |
| Solvent | 3 | 2 |
| Protein residues | 112 | 112 |
| R.m.s. deviations | | |
| Bond length (Å) | 0.003 | 0.009 |
| Bond angles (°) | 0.56 | 1.17 |
| Ramachandran plot | | |
| Favored (%) | 100.00 | 98.15 |
| Allowed (%) | 0.00 | 1.85 |
| Outliers (%) | 0.00 | 0.00 |
| Average B-factor ($Å^2$) | 87.76 | 89.43 |
| Macromolecules | 80.79 | 82.15 |
| Ligands | 75.25 | 75.52 |
| Solvent | 76.37 | 66.78 |

Statistics for the highest-resolution shell are shown in parentheses.

† a*: 2.17 Å, b*: 2.17 Å, c*: 3.01 Å
